# Tick-borne pathogens in ticks (Acari: Ixodidae) collected from various domestic and wild hosts in Corsica (France), a Mediterranean island environment

**DOI:** 10.1101/723189

**Authors:** Sébastien Grech-Angelini, Frédéric Stachurski, Muriel Vayssier-Taussat, Elodie Devillers, François Casabianca, Renaud Lancelot, Gerrit Uilenberg, Sara Moutailler

## Abstract

Corsica is a touristic mountainous French island in the north-west of the Mediterranean Sea presenting a large diversity of natural environments where many interactions between humans, domestic animals and wild fauna occur. Despite this favourable context, tick-borne pathogens (TBPs) have not systematically been investigated. In this study, a large number of TBPs were screened in ticks collected during one year from domestic and wild hosts in Corsica. More than 1,500 ticks belonging to nine species and five genera (*Rhipicephalus*, *Hyalomma*, *Dermacentor*, *Ixodes* and *Haemaphysalis*) were analysed individually or pooled (by species, gender, host and locality). A real-time microfluidic PCR was used for high-throughput screening of TBPs DNA. This advanced methodology permitted the simultaneous detection of 29 bacterial and 12 parasitic species (including *Borrelia*, *Anaplasma*, *Ehrlichia*, *Rickettsia*, *Bartonella*, *Candidatus* Neoehrlichia, *Coxiella*, *Francisella*, *Babesia* and *Theileria*). CCHF virus was investigated individually in tick species known to be vectors or carriers of this virus. In almost half of the tick pools (48%), DNA from at least one pathogen was detected and eleven species of TBPs from six genera were reported. TBPs were found in ticks from all collected hosts and were present in more than 80% of the investigated area. The detection of some pathogens DNA confirmed their previous identification in Corsica, such as *Rickettsia aeschlimannii* (23% of pools), *Rickettsia slovaca* (5%), *Anaplasma marginale* (4%) and *Theileria equi* (0.4%), but most TBPs DNA was not reported before in Corsican ticks. This included *Anaplasma phagocytophilum* (16%), *Rickettsia helvetica* (1%)*, Borrelia afzelii* (0.7%)*, Borrelia miyamotoi* (1%)*, Bartonella henselae* (2%), *Babesia bigemina* (2%) and *Babesia ovis* (0.5%). The important tick infection rate and the diversity of TBPs reported in this study highlight the probable role of animal reservoir hosts for zoonotic pathogens and human exposure to TBPs on Corsica.

## 1. INTRODUCTION

The regional sanitary importance of ticks depends on the tick species and tick-borne pathogens (TBPs) present in an area, and to a large extent on the local climate, management and breeds of livestock and human activities (Jongejan and Uilenberg, 2004). The role of ticks as vectors of human pathogens is second in importance to that of mosquitoes (Parola and Raoult, 2001) and they are worldwide the most important vectors in the veterinary field (Nicholson et al., 2009). Ticks can transmit many varieties of pathogens, including bacteria, parasites and viruses. Moreover, human tick-borne diseases are usually zoonotic and asymptomatic for non-human vertebrate hosts which most often are the reservoirs of pathogens causing human infection (Jongejan and Uilenberg, 2004).

Corsica is a French island in the western part of the Mediterranean area, situated 15 km north of Sardinia and 90 km west of Tuscany in Italy. It is the fourth Mediterranean island in size and the most mountainous and forested one. The island consists of two departments (Corse-du-Sud and Haute-Corse) and 360 communes (the smallest administrative unit in France; Fig. 1). Tourism (three million people annually, 320,000 permanent inhabitants), extensive farming (sheep, goats, pigs and cattle), hunting and hiking are important activities in Corsica (Grech-Angelini et al., 2016b). Therefore, in this context, permanent interactions occur between livestock, wildlife and humans in a small area, which certainly favour the circulation of TBPs, including zoonotic ones. Corsica is also on the route of migratory birds which create a natural link between Africa and Europe and could spread ticks infected with TBPs (Hoffman et al., 2018).

**Figure 1.**
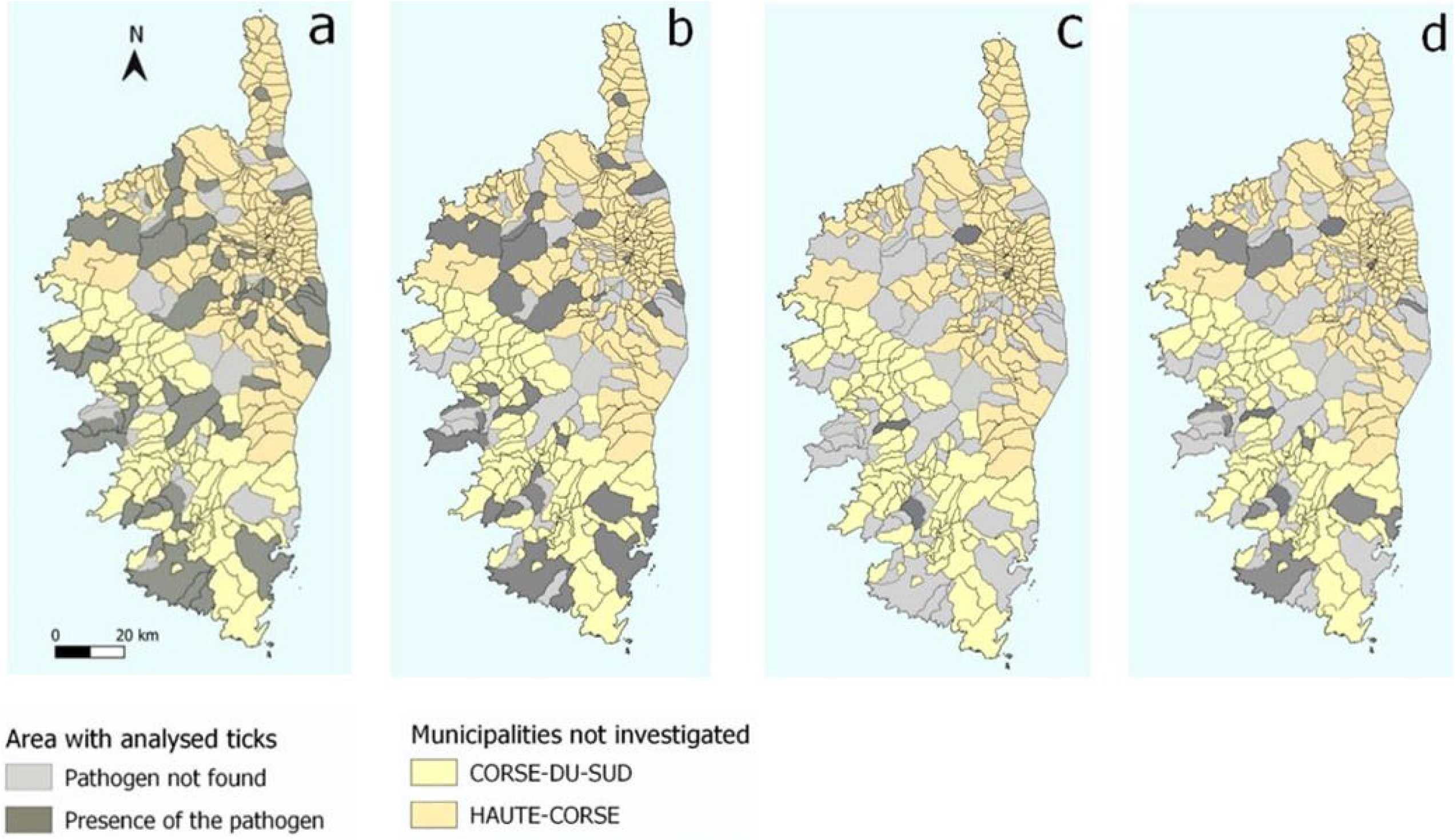
Distribution of the main DNA pathogens found in Corsican ticks. a) *Rickettsia s* pp., b) *Anaplasma* spp., c) *Borrelia* spp., d) *Babesia* spp.

Only scattered observations on the tick fauna of Corsica, mostly grouped together in a book on the ticks of France (Pérez-Eid, 2007), were available before 2014. From May 2014 to May 2015, a large tick survey (Grech-Angelini et al, 2016b) on domestic and a few wild animals led to the identification of nine species: *Rhipicephalus bursa*, *Hyalomma marginatum*, *Dermacentor marginatus*, *Rh. sanguineus* sensu lato, *Hy. scupense*, *Ixodes ricinus*, *Haemaphysalis punctata*, *Rh. (Bo.) annulatus*, and *Ha. sulcata*. The diversity of the Corsican tick species, characterized by ticks usually collected in humid environments (*I. ricinus*) and others in drier areas (*Hyalomma* spp.*, Rh. bursa*), suggested a potential high diversity of TBPs on the island.

Some of the TBPs occurring on Corsica have been reported more or less reliably in former and recent studies. Various species of the genera *Anaplasma*, *Rickettsia* and *Ehrlichia* (ICTTD, 2000; Matsumoto at al., 2004; Dahmani et al., 2017; Selmi et al., 2017; Cabezas-Cruz et al., 2019; Cicculli et al., 2019a, 2019b, 2019c), the genera *Babesia* and *Theileria* (ICTTD, 2000) have been identified, and *Borrelia burgdorferi* sensu lato was recently reported (Cicculli et al., 2019c). It remained uncertain whether all of the Corsican TBPs were known. The aim of this study was to have a large overview regarding the TBPs carried in more than 1,500 ticks collected on different Corsican animal hosts, focusing on the main pathogens of medical and veterinary importance known in the Mediterranean area, including *Borrelia* spp., *Rickettsia* spp., *Anaplasma* spp., *Francisella* spp., *Ehrlichia* spp., *Coxiella* spp., *Theileria* spp., *Babesia* spp., *Bartonella* spp., *Candidatus* Neoehrlichia and CCHF virus.

## 2. MATERIAL AND METHODS

### 2.1 Study area and tick collection

A large-scale tick collection has been conducted on domestic and a few wild animals in Corsica (Grech-Angelini et al., 2016a and 2016b). Cattle were chosen as host model because of the extensive, free-ranging livestock farming system, with a low frequency of acaricide treatments. Ticks were collected over a year (May 2014 to May 2015) in the three cattle Corsican slaughterhouses. Ticks on sheep, goats and horses were collected less systematically from May to August 2014 in three farms for each host. Ticks from domestic carnivores were provided by practising veterinarians. Ticks from wild boars, mouflons and deer were obtained respectively from hunters, from staff of the National Office for Hunting and Wildlife (ONCFS) and of the Regional Natural Park of Corsica (PNRC). Ticks were stored in 70 % ethanol at −20 °C until their identification.

Ticks were identified on their morphology; when deemed necessary, some specimens were also molecularly examined by sequencing mitochondrial *cox*1 (cytochrome *c* oxidase subunit 1) and 16S ribosomal RNA genes and ITS2 (internal transcribed spacer 2) (Grech-Angelini et al., 2016a and 2016b).

### 2.2 Pools of ticks

Tick were analysed individually or by pools consisting of two to five ticks. *Rhipicephalus bursa*, *Hy. marginatum*, *Ha. punctata*, *Rh. sanguineus* s.l. and *D. marginatus*, found in large numbers on their hosts, were grouped in pools by sex, host and locality. Other species found more rarely or with a special sanitarian interest were systematically analysed individually: *I. ricinus*, *Hy. scupense, Ha. sulcata* and *Rh*. (*Bo.*) *annulatus*.

### 2.3 DNA and RNA extraction

After washing once in 70% ethanol for 5 min and twice in distilled water for 5 min, pools of one to five ticks were crushed in 300 μl of DMEM with 10% foetal calf serum and six steel balls using the homogenizer Precellys® 24 Dual (Bertin, France) at 5500 rpm for 40s. DNA was then extracted using 100μl of the homogenate according to the Wizard genomic DNA purification kit (Promega, France) following the manufacturer’s instruction. Total DNA per sample was eluted in 50 μl of rehydration solution and stored at −20°C until further use (Michelet et al, 2014). Using 100μl of tick homogenate for *Hyalomma* spp. and *Rh. bursa* (species known to transmit or to carry CCHF virus), the Nucleospin RNA II kit (Macherey-Nagel, Duren, Germany) was used for total RNA extraction following the manufacturer’s instruction (Gondard et al., 2018).

### 2.4 Detection of tick-borne pathogen DNA or RNA

#### Bacteria and parasites

Forty-one sets of primers and probes were used in this study to detect TBPs (29 bacterial and 12 parasitic species, Tab. 1). The BioMark™ real-time PCR system (Fluidigm, USA) was used for high-throughput microfluidic real-time PCR amplification using 48.48 dynamic arrays (Fluidigm). These chips dispense 48 PCR mixes and 48 samples into individual wells, after which on-chip microfluidics assemble PCR reactions in individual chambers prior to thermal cycling resulting in 2,304 individual reactions (Michelet et al., 2014).

**Table 1.**
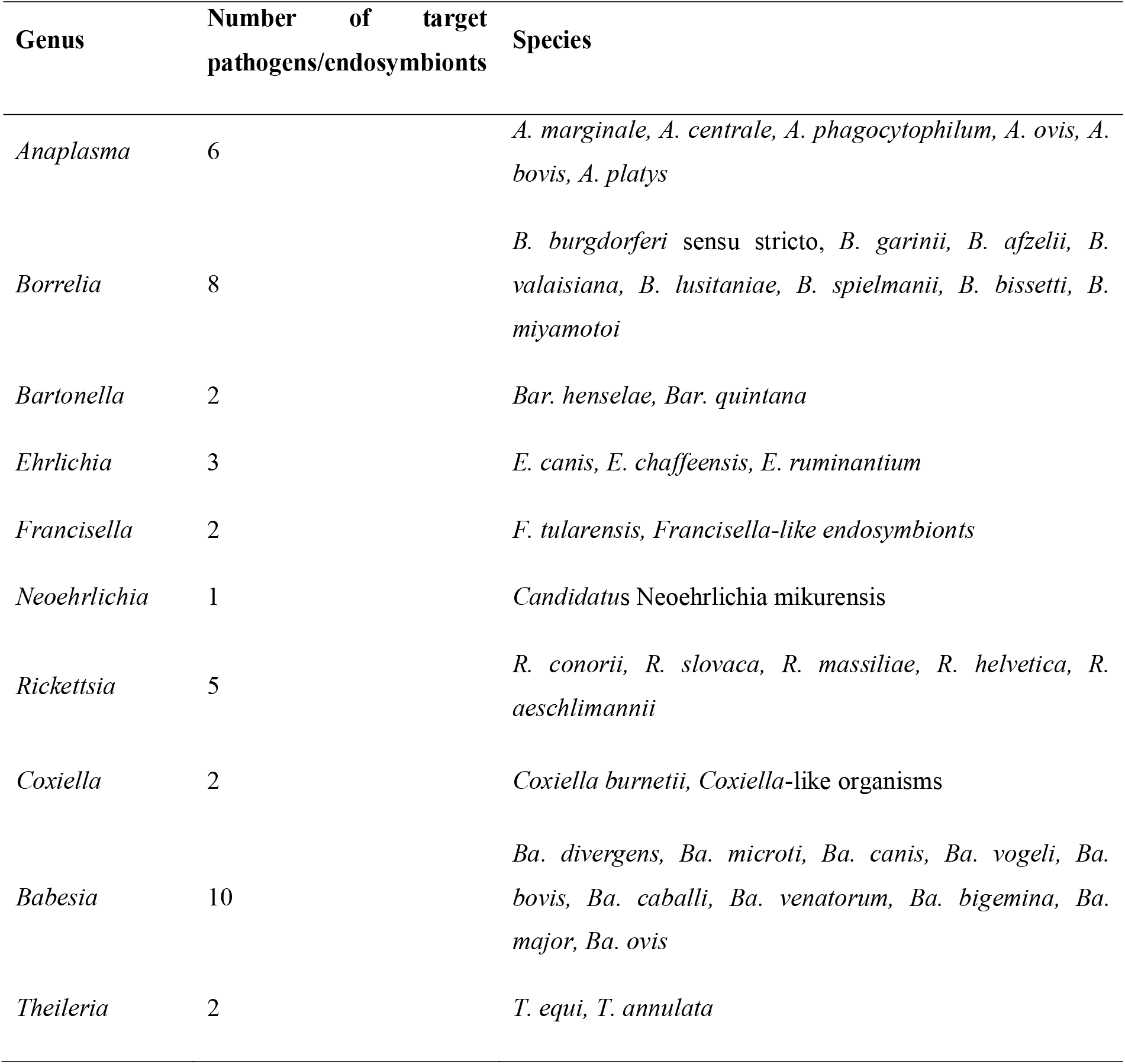
List of pathogens (and endosymbionts) detectable by the real-time PCR chip (Biomark™)

Conventional PCR using primers targeting genes or regions other than those of the BioMark™ system were used to confirm the presence of pathogenic DNA in the field samples. Amplicons were sequenced by Eurofins MWG Operon (Germany), and then assembled using BioEdit software (Ibis Biosciences, Carlsbad). An online BLAST (National Center for Biotechnology Information) was used to compare results with published sequences listed in GenBank sequence databases (Michelet et al, 2014; Gondard et al., 2019).

#### Specific detections of CCHF virus and *Theileria* spp

CCHF virus RNA was specifically investigated in tick species known to transmit, or at least able to carry it: *Hy. marginatum, Hy. scupense*, and *Rh. bursa*. The presence of *Theileria* spp. DNA was also investigated by specific real-time PCR in *Hy. scupense* and *Haemaphysalis* spp. because *Hy. scupense* is an efficient vector of *T. annulata* and because *T. buffeli*, of which *Ha. punctata* is a vector, has already been reported in Corsica (ICTTD, 2000) (for primers and probes see Gondard et al., 2018 and 2019).

### 2.5 Data analysis

The infection rate is given for all pathogens in each tick species and each host. The infection rate of individually analysed tick species is the real one detected. The infection rate of pooled tick species (cf. Methods, 2.4) means that in an infected pool there was at least one tick that gave a positive result. To assess differences in tick species infection between development stage and sex a Kruskal-Wallis test was used. Differences were considered as statistically significant if *p* < 0.05.

## 3. RESULTS

### 3.1 Collected ticks and pool preparation

More than three thousand ticks (3,134) from nine species and five genera were collected from May 2014 to May 2015 (Grech-Angelini et al., 2016b). Almost half of them (1,523) were analysed to detect animal and human pathogens DNA or RNA (Tab. 2). A total of 569 samples were analysed (consisting of one to five ticks): 269 adult females, 261 adult males and 39 nymphs. Nymphs of four species were found: *Rh. bursa*, *Hy. scupense, Rh. sanguineus* group and *Rh. (Bo.) annulatus* (Tab. 2). *Rhipicephalus bursa* ticks (4.8 ticks per pool on average), *Hy. marginatum* (4.1), *Dermacentor marginatus* (4.1)*, Rh. sanguineus* sensu lato (3.7) and *Ha. punctata* (2.5) were systematically pooled whereas ticks from the other four species were analysed individually (*Rh. (Bo.) annulatus*, *Ha. sulcata, Hy. scupense* and *Ixodes ricinus*).

**Table 2.**
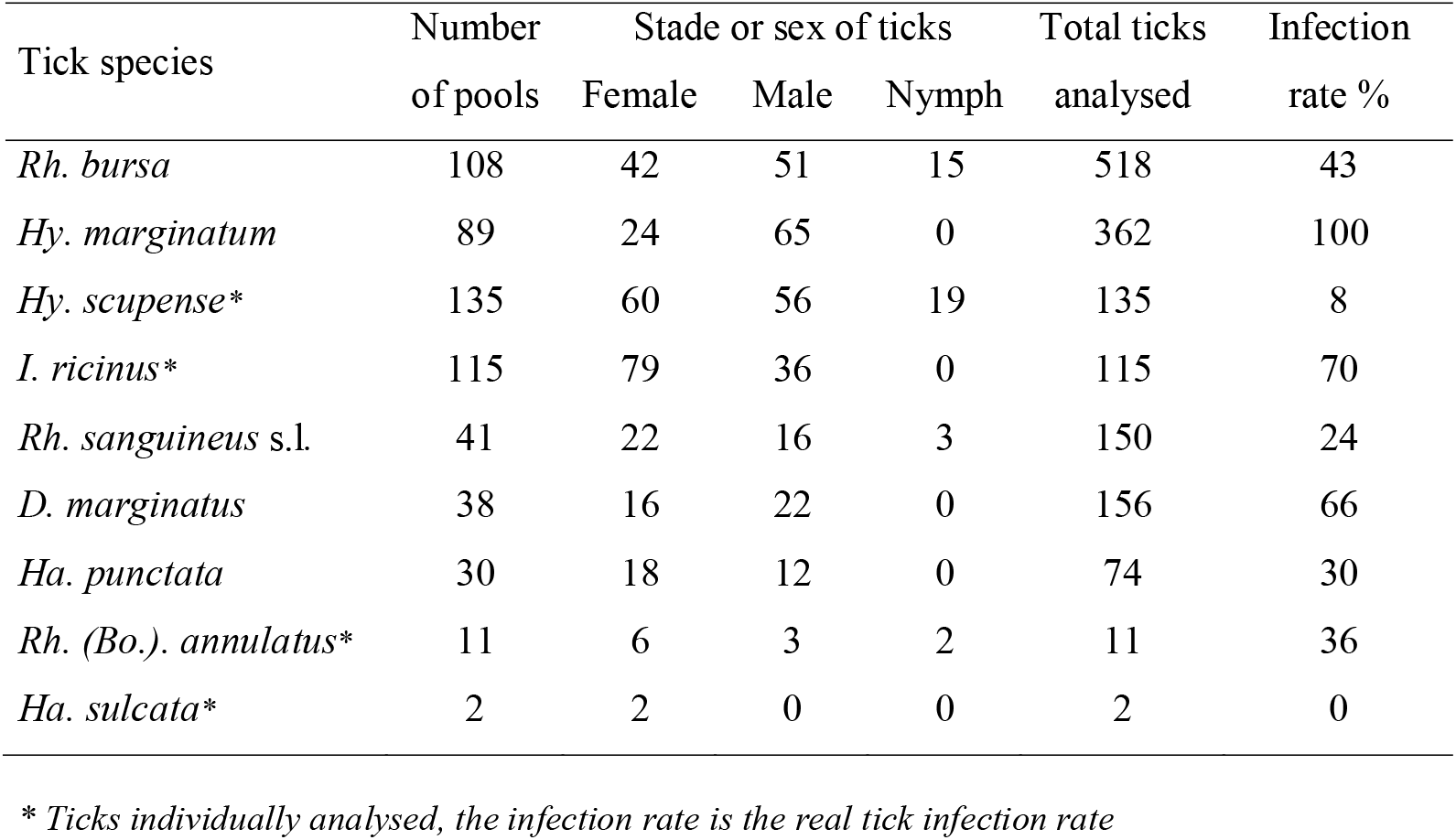
Infection rates of individual ticks or pools of ticks collected in Corsica.

| Tick species | Number<br>of pools | Stade or sex of ticks |  |  | Total ticks<br>analysed | Infection<br>rate % |
| --- | --- | --- | --- | --- | --- | --- |
|  |  | Female | Male | Nymph |  |  |
| *Rh. bursa* | 108 | 42 | 51 | 15 | 518 | 43 |
| *Hy. marginatum* | 89 | 24 | 65 | 0 | 362 | 100 |
| *Hy. scupense* * | 135 | 60 | 56 | 19 | 135 | 8 |
| *I. ricinus* * | 115 | 79 | 36 | 0 | 115 | 70 |
| *Rh. sanguineus* s.l. | 41 | 22 | 16 | 3 | 150 | 24 |
| *D. marginatus* | 38 | 16 | 22 | 0 | 156 | 66 |
| *Ha. punctata* | 30 | 18 | 12 | 0 | 74 | 30 |
| *Rh. (Bo.). annulatus* * | 11 | 6 | 3 | 2 | 11 | 36 |
| *Ha. sulcata* * | 2 | 2 | 0 | 0 | 2 | 0 |

\* Ticks individually analysed, the infection rate is the real tick infection rate

### 3.2 Detection of TBPs DNA or RNA in tick species

As for each tick species, the proportion of infected pools was not significantly different between development stage and sex (*p* > 0,05), the identified pathogens were presented by tick species, not separately for the stage of development and the sex of adults. Almost half of the samples (48%) were positive for at least one pathogen DNA (Tab. 2) and among them 12% were positive for two pathogens DNA or more. The most infected tick species were *Hy. marginatum* (100% of pools), *I. ricinus* (70% of the individual ticks) and *D. marginatus* (66% of pools). Pathogen DNA was found in all tick species collected in Corsica except in *Ha. sulcata*, but only two specimens were analysed (Tab. 2 and Tab. 3).

**Table 3.** Proportion (%) of infected pools (or ticks)

| Tick species<br>(pools; ticks analysed) |  |  |  |  |  |  |
| --- | --- | --- | --- | --- | --- | --- |
|  | *Borrelia* | *Rickettsia* | *Anaplasma* | *Babesia* | *Bartonella* | *Theileria* |
| *Rh. bursa* (108;518) | 0 | 27 | 15 | 12 | 5 | 1 |
| *Hy. marginatum* (89;362) | 0 | 100 | 7 | 0 | 1 | 0 |
| *Hy. scupense* (135;135)* | 0 | 6 | 2 | 0 | 0 | 0 |
| *I. ricinus* (115; 115)* | 5 | 6 | 64 | 0 | 2 | 0 |
| *Ha. punctata* (30;74) | 13 | 0 | 13 | 0 | 3 | 0 |
| *Ha. sulcata* (2;2)* | 0 | 0 | 0 | 0 | 0 | 0 |
| *Rh. sanguineus* s.l. (41;150) | 0 | 17 | 12 | 0 | 2 | 0 |
| *Rh. (B.) annulatus* (11;11)* | 0 | 0 | 18 | 0 | 9 | 9 |
| *D. marginatus* (38;156) | 0 | 66 | 3 | 0 | 3 | 0 |

\*Ticks individually analysed.

### 3.3 Pathogens identified

DNA of eleven pathogen species from six genera were identified in the ticks: *Borrelia* spp., *Rickettsia* spp., *Anaplasma* spp., *Babesia* spp., *Bartonella* spp. and *Theileria* spp.

#### *Borrelia* spp

*Borrelia* spp. DNA was detected in two tick species and two different *Borrelia* were identified. DNA of *Borrelia miyamotoi* was found in *Ixodes ricinus* (2% of individual ticks) and in *Ha. punctata* (13% of the pools). A sequence was obtained from *I. ricinus* (GenBank, accession number: MK732472), and showed 100% homology with reference sequences from *B. miyamotoi* strain isolated in Austria (AN: KP202177). DNA of *B. afzelii* was reported only in *I. ricinus* (4% of ticks). Unfortunately, the sequence could not be obtained.

#### *Rickettsia* spp

DNA of *Rickettsia* spp. was detected in six tick species (Tab. 3). Three species of *Rickettsia* were identified. *Rickettsia aeschlimannii* was found in *Rh. bursa* (27% of pools)*, Hy. marginatum* (100% of pools)*, Rh. sanguineus* s.l. (10% of pools) and *Hy. scupense* (5% of individual ticks). The four sequences obtained from *Hy. marginatum* and *Hy. scupens*e presented 100% identity. One sequence was deposited in GenBank (AN: MK732478), and showed 99% homology with reference sequences from *R. aeschlimannii* strain isolated in Egypt (HQ335153). *Rickettsia slovaca* was mostly identified in *D. marginatus* (66% of pools). Three pools of *Rh. sanguineus* s.l. (7%) and one *Hy. scupense* tick (0.7%) were also positive for this bacterium. The sequence obtained from *Hy. scupense* (GenBank, AN: MK732480) showed 99% homology with reference sequences from *R. slovaca* strain isolated in United States (AN: KR559551). *Rickettsia helvetica* was detected only in *I. ricinus* (6% of ticks) but the sequence could not be obtained.

#### *Anaplasma* spp

DNA of *Anaplasma* spp. was detected in eight tick species (Tab. 3) and two species were recorded: *A. phagocytophilum* and *A. marginale*. *Anaplasma phagocytophilum* DNA was mainly found in *I. ricinus* (63% of ticks), but it was also reported in *Rh. sanguineus* s.l. (7% of pools), *Rh. bursa* (7% of pools), *Hy. marginatum* (1% of pools), *Hy. scupense* (2% of ticks), *D. marginatus* (3% of pools) and *Ha. punctata* (7%). Unfortunately, the sequence of *A. phagocytophilum* could not be obtained. *Anaplasma marginale* DNA was frequently detected in *Rh. (Bo.) annulatus* (18% of ticks) but it was also identified in *Rh. bursa* (8% of pools), *Hy. marginatum* (6% of pools), *I. ricinus* (2% of ticks), *Ha. puncta* (7% of pools) and *Rh. sanguineus* s.l. (7% of pools). The sequence obtained from *Rh. (Bo.) annulatus* (GenBank, AN: MK732471) showed 99% homology with reference sequences from *A. marginale* strain isolated in South Africa (AN: AF414873).

#### *Babesia* spp

DNA of *Babesia* spp. was detected only in *Rh. bursa* and two species were identified. *Babesia bigemina* was found in 9% of the pools of *Rh. bursa.* The two sequences obtained presented 100% identity. One sequence was deposited in GenBank (AN: MK732475) and showed 100% homology with reference sequences from *Ba. bigemina* strain from Virgin Islands (AN: EF458206). *Babesia ovis* was identified in 3% of *Rh. bursa* pools. The sequence obtained (GenBank, AN: MK732477) showed 92% homology with reference sequences from *Ba. ovis* strain isolated in Sudan (AN: AY260171).

#### *Bartonella* spp

*Bartonella henselae* was the only species of the genus *Bartonella* detected and its DNA was identified in seven tick species (Tab. 3) but mainly in *Rh. bursa* (5% of pools), *I. ricinus* (3% of ticks), *Rh. (Bo.) annulatus* (9% of ticks) and *Ha. punctata* (3% of pools). The five sequences obtained in *Rh. bursa* and *I. ricinus* presented 100% identity. One sequence was deposited in GenBank (AN: MK732473) and showed 100% homology with reference sequences from *Bar. henselae* strain Houston-1 (AN: BX897699).

#### *Theileria* spp

*Theileria equi* DNA was reported in one pool of *Rh. bursa* (1%) and in one *Rh. (Bo.) annulatus* (9% of ticks) (Tab. 3). The sequence obtained from *Rh. bursa* (GenBank, AN: MK732476) showed 99% homology with reference sequences from *T. equi* isolated in Brazil (AN: KJ573370). No *Theileria annulata* DNA was detected by the BioMark™ real-time PCR system in the analysed ticks. Individual PCR for *Theileria* spp. did not show the presence of any other species of *Theileria* in the tick species analysed (*Hy. scupense* and *Haemaphysalis* spp.).

#### Others pathogens and micro-organisms

*Francisella tularensis* was not identified in this study but *Francisella-*like endosymbionts were detected in five tick species: *Hy. marginatum* (90% of pools), *Hy. scupense* (7% of ticks), *I. ricinus* (4% of ticks), *Rh. sanguineus* s.l. (10% of pools) and *Rh. (Bo.) annulatus* (27% of ticks). Neither DNA of *Coxiella* spp. nor DNA of *Candidatus* Neoehrlichia mikurensis were identified in this study. RNA of CCHF virus was not found in *Hy. marginatum* (89 pools; 362 ticks), *Rh. bursa* (108; 518) and *Hy. scupense* (135 ticks).

For some *Rickettsiae*, the exact species could not be determined t. These *Rickettsiae* were reported in seven tick species: *Rh. bursa* (8% of pools), *Hy. scupense* (2% of ticks), *I. ricinus* (9% of ticks), *Rh. sanguineus* s.l. (12% of pools), *D. marginatus* (11% of pools), *Ha. punctata* (3% of pools) and *Rh. (Bo.) annulatus* (9% of ticks). One sequence was obtained from *Rh. bursa* (GenBank, AN: MK732479) and showed 99% homology with reference sequences from *Rickettsia*-like endosymbiont strain 162 citrate synthase (gltA) gene (AN: JQ925589).

### 3.4 Presence of pathogens in ticks collected on the different hosts

Ticks were collected from different hosts: 404 samples (consisting to one to five ticks) originated from cattle, 19 from goats, 12 from sheep, 24 from horses, 22 from dogs, four from cats (Tab. 4), 41 from wild boars, 26 from mouflons, 13 from deer and two from hedgehog and birds (Tab. 5). For birds and deer, as the collected ticks were all *I. ricinus*, they were individually analysed.

**Table 4.** Pathogen DNA detected in ticks collected from domestic animals.

| Host | Analysed<br>pools (ticks) | Tick species pools (or individual ticks*) | % infected<br>pools (or ticks*) | Pathogens identified<br>(% of infected pools or ticks*) |
| --- | --- | --- | --- | --- |
| Cattle | 404 (894) | *Rh. bursa* (54), *Hy. marginatum* (67),<br>*Hy. scupense* (135), *I. ricinus* (98), *Rh.*<br>*sanguineus* s.l. (9), *D. marginatus* (4),<br>*Ha. punctata* (26), *Rh. (B.) annulatus*<br>(11) | 47 | *Borrelia* spp. (3%), *Rickettsia* spp.<br>(25%), *Anaplasma* spp. (22%),<br>*Babesia* spp. (3%), *Bartonella* spp.<br>(2%), *Theileria* spp. (0.3%) |
| Goat | 19 (91) | *Rh. bursa* (18), *Ha. punctata* (1) | 47 | *Rickettsia* spp. (37%), *Anaplasma*<br>spp. (2%), *Babesia* spp. (5%),<br>*Bartonella* spp. (11%) |
| Sheep | 12 (45) | *Rh. bursa* (9) *Hy. marginatum* (2), *Rh.*<br>*sanguineus* s.l. (1) | 58 | *Rickettsia* spp. (58%) |
| Dogs | 22 (86) | *Hy. marginatum* (1), *Rh. sanguineus* s.l.<br>(21) | 14 | *Rickettsia* spp. (9%), *Anaplasma*<br>spp. (4%), |
| Horses | 24 (106) | *Rh. bursa* (6), *Hy. marginatum* (16), *Rh.*<br>*sanguineus* s.l. (1),<br>*D. marginatus* (1), | 83 | *Rickettsia* spp. (79%), *Bartonella*<br>spp. (4%), *Theileria* spp. (4%) |
| Cats | 4 (6) | *I. ricinus* (1), *Rh. sanguineus* s.l. (3) | 25 | *Anaplasma* spp. (25%) |

**Table 5.**
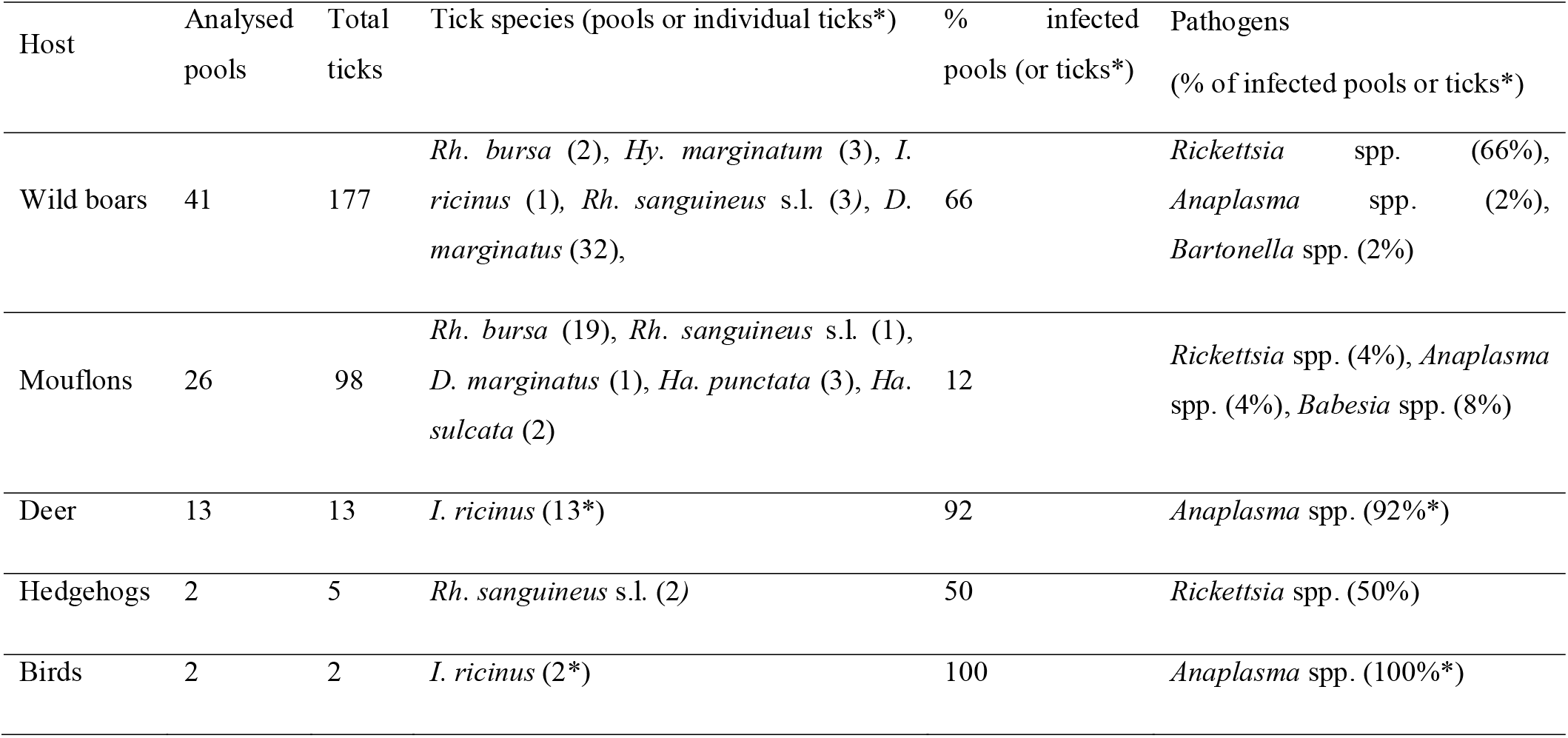
Pathogen DNA detected in ticks collected from wild animals.

*Borrelia* spp. DNA was found only in ticks from cattle; *Borrelia miyamotoi* and *Bo. afzelii* infected respectively 2% and 1% of pools from this host.

*Rickettsia* spp. DNA was found in ticks from almost all collected hosts. *Rickettsia aeschlimannii* was the most important pathogen infecting ticks from domestic ruminants and horses, reported in 23% of pools from cattle, 37% from goats, 58% from sheep and 75% from horses. It was rarely found in pools from dogs (5%), wild boars (7%) and mouflons (4%). *Rickettsia slovaca* DNA was mainly found in pools from wild boars (59%) but it was also reported in pools from cattle (0.5%), horses (4%), dogs (5%) and hedgehog (one positive pool). *Rickettsia helvetica* DNA was identified only in ticks collected from cattle (2%).

*Anaplasma phagocytophilum* DNA was reported in ticks from cattle (18% of pools), goat (11% of pools), cats (25% of pools), wild boars (2% of pools), deer (92% of individual ticks) and birds (two positive ticks) whereas *A. marginale* DNA was found in pools from cattle (6%) and mouflons (4%).

*Babesia bigemina* DNA was detected mainly in ticks sampled on cattle (2% of pools) but one pool of ticks from goats was also positive for this pathogen. *Babesia ovis* DNA was reported in two hosts, one pool of ticks collected on cattle and two pools from mouflons were positive.

*Bartonella henselae* DNA was reported in pools from cattle (2%), goat (11%), horses (4%) and wild boars (2%). *Theileria equi* DNA was found in one pool from cattle and one from wild boars.

### 3.5 Geographical distribution of TBPs found in ticks

Ticks from 82 municipalities were analysed and among them, pathogen DNA was found in 68 (Fig. 1). *Rickettsia* spp. were widespread (Fig. 1a) as they were reported in more than 80% of the municipalities investigated. *Rickettsia slovaca* DNA was found in ticks collected from 13 municipalities, *R. aeschlimannii* DNA from 49 and *R. helvetica* DNA was reported from a single municipality in the centre of Corsica (Haute-Corse). *Anaplasma* spp. (Fig. 1b) were also distributed in a large part of the investigated area, found in 37% of the municipalities. *Anaplasma phagocytophilum* DNA was reported in 19 municipalities and *A. marginale* DNA in 16. *Borrelia* spp. were found in five municipalities (Fig. 1c). *Borrelia miyamotoi* DNA was identified in all these municipalities whereas *B. afzelii* DNA was reported only in one from Haute-Corse. *Babesia* spp. were identified in 10 municipalities as well in Corse-du-Sud as in Haute-Corse (Fig. 1d). *Bartonella henselae* DNA was identified in 12 municipalities whereas *T. equi* DNA was found in two municipalities near the eastern coast in Haute-Corse (results not shown).

## 4. DISCUSSION

In this study, a method using multiple primers/probe sets was implemented to perform high-throughput detection of TBPs DNA. This large-scale investigation has (i) enabled the detection of rare pathogens, (ii) generated prevalence estimations for pathogens, thus creating a comprehensive overview of the epidemiological situation for 29 bacteria and 12 parasites present or not found in ticks from Corsica. A positive tick or a positive pool means it contains DNA (or RNA) sequences similar to those of corresponding genes of TBPs, and does not necessarily mean that the TBP is present in the tick. For the sake of commodity, the term “infection rate” is often used in this study but it means more “positive rate for pathogen DNA”. As all collected ticks were removed from their hosts, potentially infected by pathogens, the identification of pathogen DNA in a tick suggests its presence in Corsica but the tick species is not necessarily a biological vector of the pathogen. The micro-organism or its DNA may be simply present in the ingested blood meal, the host having been infected by a real vector species of tick (Estrada-Peña et al., 2013). However, it has to be noted that in this survey, the main pathogens were most often identified in their natural vector suggesting a higher link between one pathogen and its natural vector than with other tick species. Almost 70% of the *R. aeschlimannii* were detected in *Hy. marginatum*, 86% of the *R. slovaca* in *D. marginatus*, 80% of *A. phagocytophilum* in *I. ricinus*, more than 60% of *A. marginale* in ticks from the genus *Rhipicephalus* and all the *Babesia* spp. were identified in *Rh. bursa*.

### 4.1 Tick-borne pathogens reported for the first time in Corsican ticks

In almost half of the pools, DNA of at least one pathogen was reported and 11 species of pathogens from six genera were identified. Among them, seven were reported for first time in Corsican ticks: *B. afzelii*, *B. miyamotoi*, *R. helvetica*, *A. phagocytophilum*, *Ba. ovis*, *Ba. bigemina* and *Bar. henselae*.

*Borrelia afzelii* DNA, found in 4% of the Corsican *I. ricinus*, belongs to the *Borrelia burgdorferi* s.l. group, the causative agents of Lyme disease, the most important zoonotic infection in Europe. The presence of *B. burgdorferi* s.l. is linked to its natural vectors, ticks of the genus *Ixodes*, mainly *I. ricinus* in the Mediterranean area. This pathogenic agent was believed to be rare or even absent from Corsica, but a few cases of Lyme disease have been reported in recent years by the French National Centre of *Borrelia* and Santé Publique France (Vandenesch et al., 2014). *Borrelia burgdorferi* s.l. was reported in Continental France, continental Italy, Spain (Habálek *et al*., 1997) and also in northern Africa (Tunisia, Algeria and Morocco; Benredjem et al., 2014) but has not been found in ticks from Sardinia and Sicilia, two Mediterranean islands with a drier climate than Corsica. Recently, it was reported in *I. ricinus* collected from rats in Corsica (Cicculli et al., 2019c), but without determining the exact species. The identification of *B. afzelii* DNA in ticks confirmed the exposure of Corsican population to Lyme disease.

DNA of *Borrelia miyamotoi*, first identified in 1995 in ticks from Japan (Fukunaga et al., 1995), was also detected for the first time in Corsica. It belongs to the relapsing fever group of *Borrelia* and its natural vectors are *Ixode*s ticks. *Borrelia miyamotoi* is currently considered as an emerging pathogen affecting humans. Relapsing fever borreliosis is characterized by influenza-like illness and one or more relapse episodes of bacteraemia and fever (Platonov *et al*., 2011). *Borrelia miyamotoi* was reported in 3% of ticks from continental France (Cosson et al., 2014) and 0.7% of northern Italy (Ravagean *et al*., 2018). Although no human case of *B. miyamotoi* was identified in Corsica and in the Mediterranean area, the bacterium has been found in humans throughout Europe (Siński et al., 2016).

*Rickettsia helvetica* DNA was only found in *I. ricinus* which is a natural vector of this bacterium. It was initially reported in 1979 in Switzerland from *I. ricinus* (Beati et al., 1993) and now the species has been identified in ticks from many countries over the world. It is especially well spread in the Mediterranean area including continental France (Parola *et al.*, 1998), Italy (Beninati *et al.*, 2002), Spain (Fernández-Soto *et al.*, 2004), Croatia (Dobec *et al*., 2009), Algeria (Kernif *et al.*, 2012) and Tunisia (Sfar *et al.*, 2008). Human cases of rickettsiosis caused by *R. helvetica* have been reported in Italy and continental France (Portillo *et al.*, 2015). *Rickettsia helvetica* is a member of the Spotted Fever Group (SFG) and reportedly causes a self-limiting illness associated with headache and myalgias (Parola et al., 2013).

*Anaplasma phagocytophilum* DNA was mostly found in *I. ricinus*, an important vector of this zoonotic agent responsible for human granulocytic anaplasmosis (HGA), tick-borne fever (or pasture fever) of ruminants and equine anaplasmosis. In Corsica, this pathogen was relatively unknown, and was previously only detected in a single sample of bovine blood (ICTTD, 2000). There is no HGA case reported in Corsica, but in continental France the first one was identified in 2003 and other were infrequently reported (Edouard *et al*., 2012). Human cases occurred also in continental Italy (Ruscio and Cinco, 2003). A recent study showed that, in the French Basque Country, 22.4% of collected ticks contained *A. phagocytophilum* DNA (Dahmani *et al*., 2017). It was also reported from Sardinia (Alberti *et a*l., 2005), Sicily (Torina *et al*., 2010) and Northern Africa (Dahmani *et al*., 2015).

*Babesia* spp. are protozoan blood parasites with more than 100 described species. In this study, *Ba. bigemina* and *Ba. ovis* DNA were found for the first time in ticks collected in Corsica. *Babesia bigemina* is a causal agent for bovine babesiosis and it occurs in most areas of the world (Uilenberg, 2006)*. Babesia ovis* is a causative agent of ovine babesiosis, with strains varying in pathogenicity and is also widespread throughout the world (Uilenberg, 2006).

Ticks (especially *I. ricinus)* are highly suspected to be among the vectors of *Bartonella* species. *Bartonella henselae* DNA was reported for the first time in ticks from Corsica and was found in seven tick species. It causes an infection commonly encountered in cats (cat scratch disease) and potentially dogs and humans worldwide (Álvarez-Fernández et al., 2018). In Sardinia, it was found in at least 0.2% of the collected ticks (Chisu et al., 2018).

### 4.2 Other TBPs identified in Corsica

*Rickettsia* species were the pathogens with the highest infection rate found in the Corsican pools. They are known as zoonotic agents and, if their pathogenicity and the reservoir role for animals are discussed, most species of this genus cause important human diseases (Davoust et al., 2010). The real impact of rickettsial diseases in Corsica is unknown, but some cases of Mediterranean spotted fever (MSF, caused by *R. conorii*) are reported by local medical doctors and the French National Reference Centre (CNR) for *Rickettsia* species. A former seroepidemiological study showed in 1985 that 4.8% of people were exposed to theses pathogens in Corse-du-Sud (Raoult et al., 1985). *Rickettsia aeschlimannii* DNA was the most frequently detected, infecting 100% of *Hy. Marginatum* pools, one of its main natural vectors (Matsumoto *et al*, 2004), confirming the important presence of this bacterium in Corsica, already reported in 74% of *Hy. marginatum* collected by Matsumoto *et al*. (2004). *Rickettsia aeschlimannii* was first isolated from *Hy. marginatum* in 1997 from Morocco (Beati *et al*., 1997) and is now reported in the whole Mediterranean area (Parola et al., 2013). *Rickettsia aeschlimannii* infections in humans cause spotted fever; this has previously been confirmed in northern Africa and South Africa, and in 2010 a first case occurred in Southern Europe, in a Greek patient (Germanakis et al., 2013). Given the important infestation rate of this pathogen in Corsican ticks, human exposure to *R. aeschlimannii* infection is high and human cases of tick-borne spotted fever acquired in Corsica could often be due to *R. aeschlimannii*.

*Rickettsia slovaca* DNA was identified mainly in *D. marginatus* which is an important natural vector of this bacterium. This report confirmed the results of Selmi et al. (2017) who identified *R. slovaca* in *D. marginatus* collected on Corsican vegetation. *Rickettsia slovaca* was first isolated in 1968 in a *D. marginatus* collected in Slovakia and is largely spread in the Mediterranean area. It was found in Sardinia and continental Italy (Chisu et al., 2016 and Selmi et al, 2017), continental France (Michelet et al., 2017), and Spain (Fernández-Soto et al., 2006). The first proven human case of *R*. *slovaca* infection was reported in 1997 in continental France and this micro-organism is now known as a cause of disease in various European Mediterranean countries (de Sousa et al., 2013). It is associated with TIBOLA (tickborne lymphadenopathy) syndrome, characterized by lymph node enlargement and scalp eschars (Parola et al., 2013). So far, the TIBOLA syndrome has not been reported in Corsica. This study did not allow to identify DNA of other *Rickettsia* spp. previously reported in Corsica as *Rickettsia felis*, a SFG *Rickettsia*, found in a flea (*Archaeopsylla erinacei*) collected from a fox (Marié *et al*., 2012). Matsumoto *et al*. (2005) reported the species *R. massiliae* in *Rh. sanguineus* s.l. collected on dogs in Corse-du-Sud and a recent study identified *R. africae* DNA in an *Amblyomma variegatum*, tick species not established in Corsica but whose one adult was collected, certainly following introduction of a nymph by migrating bird (Cicculli et al., 2019a). DNA of another species of *Rickettsia*, Candidatus *Ri. barbariae*, was recently detected in ticks collected from Corsican cattle (Cicculli et al., 2019c), but its pathogenicity remains unknown.

This study also showed that *Anaplasma* species are widespread in Corsican ticks. *Anaplasma marginale* DNA was found in ticks from the genus *Rhipicephalus* which are its natural vectors but also in *Hy. marginatum, I. ricinus* and *Ha. punctata*. *Anaplasma marginale* DNA was formerly reported in blood from Corsican cattle (ICTTD, 2000) and two recent studies detected it in *Rh. bursa* collected on cattle confirming its presence on the island (Dahmani et al., 2017 and Cicculli et al., 2019c). *Anaplasma marginale* is distributed worldwide in tropical and subtropical regions. It is the causative agent of erythrocytic bovine anaplasmosis that can affect various species of domestic and wild ruminants (Aubry and Geale, 2011). From islands near to Corsica, it was identified in Sardinia (Zobba et al, 2014) and Sicily (Torina et., 2010), but there is no report of its presence in continental France. *Anaplasma ovis* DNA, not found in this study, was recently reported in blood (52%) and ticks (*Rh. bursa*) collected from Corsican goats (Cabezas-Cruz et al., 2019). *Anaplasma bovis* and *A. omatjenne* were formerly reported in cattle blood (ICTTD, 2000) and other potential new species of *Anaplasma* were described from Corsican sheep blood (Dahmani et al., 2017).

Concerning *Babesia* species, *B. bovis*, not found in this study, was reported earlier from Corsican cattle blood (ICTTD, 2000). *Babesia bovis* occurs in most subtropical and tropical regions of the world (Uilenberg, 2006), and two of its proven vectors occur on Corsica (*Rh. bursa* and *Rh. (B.) annulatus*). Bovine babesiosis affecting cattle in Corsica is probably due either to *B. bigemina* or to *B. bovis*.

No species from the genus *Ehrlichia* were found in this study although *E. minacensis*, a species unknown so far from the Mediterranean area, has been detected recently in a *Hy. marginatum* (Cicculli et al., 2019b).

*Theileria equi* DNA was identified in two pools. It is a causative agent of equine piroplasmosis, that can be also due to *Babesia caballi* (not detected in this study). As the disease is largely spread in the island, it is well known by local veterinary practitioners and horse owners. *Theileria annulata* DNA has not been reported so far in the Corsican tick population. The high occurrence in Corsica of *Hy. scupense*, one of its main natural vectors, highlights however the risk of transmission of *T. annulata* to the local cattle population (Grech-Angelini et al., 2016a). *Theileria buffeli*, a benign cattle pathogen, already detected in Corsica (ICTTD, 2000), was not found in this study.

A total of 332 pools (1,015 ticks) of three species (*Hy. marginatum*, *Hy. scupense* and *Rh. bursa*) were tested for CCHF virus RNA and all were negative. The CCHF virus is mainly transmitted by ticks of the genus *Hyalomma* and *Hy. marginatum* is its main natural vector in Europe. Crimean-Congo haemorrhagic fever is now the most worldwide tick-borne viral infection of humans, and it can lead to haemorrhagic manifestations and considerable mortality. It is also an important emerging zoonotic disease in Turkey and south-eastern Europe (Dreshaj et al., 2016). Moreover, in 2016, the first autochthonous human cases have been reported in Spain showing that the disease can occur in western Europe (Negredo et al., 2017). Lately, CCHF RNA has been reported in a nymph of *Hy. rufipes* on a migrating bird on the small Italian island of Ventotene, identifying migrating birds as possible introduction pathway for CCHF virus into western Europe (Mancuso *et al*., 2019). *Hyalomma rufipes*, an important natural vector of CCHF virus in subSaharan Africa, was ever found on migrating birds in Corsica (Pérez-Eid, 2007). It is possible that CCHF virus was not detected in this study due to an insufficient sample size of analysed ticks or a deterioration of the genetic material. Because of the high frequency of *Hy. marginatum* in Corsica (the second most abundant tick species collected on animal hosts, Grech-Angelini et al., 2016b), it would be necessary to closely monitor the possible emergence of CCHF virus. More ticks, mainly *Hy. marginatum*, should be analysed and animal sera should be collected to test the presence of antibodies against CCHF virus in domestic ruminants which are known to be asymptomatic reservoir of CCHF virus (Bente et al., 2013).

### 4.3 Potential Corsican animal reservoirs for zoonotic pathogens and human exposure

These data concerning the presence of TBPs DNA on Corsica Island will be useful for studying possible future emerging diseases and the role of animals as reservoirs of TBPs. This study demonstrated the presence of expected pathogens DNA, as well as unexpected pathogens. Among the eleven pathogens reported in ticks collected from animal hosts, seven are zoonotic: *B. miyamotoi, B. afzelii, R. aeschlimannii, R. slovaca, R. helvetica, A. phagocytophilum* and *Bar. henselae*. Most of them are asymptomatic, or their effects on animal health are unknown. *Rickettsia aeschlimannii* DNA was reported in a high proportion in ticks from domestic ruminants and horses whereas *R. slovaca* DNA was mostly identified in ticks from wild boars and *R. helvetica* DNA only in ticks from cattle. All positive pools for *Borrelia* spp. consisted of ticks collected from cattle. These domestic or wild animals could be specific reservoirs for the respective cited pathogens. *Anaplasma phagocytophilum* was mainly found in *I. ricinus* on various hosts including migrating birds that showed the potential to import pathogens on Corsica by this pathway. Even if no human cases of these pathogens have been diagnosed in Corsica to date, the important interactions that occur between domestic animals, wild fauna and humans highlight the risk for the human population. Some of these zoonotic pathogens could circulate undetected in the Corsican human population. Such studies may help avert that more efforts should be devoted to the surveillance of the animal reservoirs and the human exposure. Pathogen DNA is not enough to be able to confirm the presence and the circulation of those pathogens in Corsica. Nevertheless, this first step will allow to perform in-depth research on those interesting pathogens to better characterize their epidemiological cycle. Indeed, serological survey in domestic and wild animals and humans need to be carry out, as well as trying to isolate those pathogens from infected ticks and/or infected animals to be able to conclude regarding the risk for human and animal health.

## 5. CONCLUSION

In this study, 569 tick pools (1,523 ticks) collected from animals on Corsica were analysed to investigate the presence of 27 bacteria and 12 parasites by a high throughput real-time microfluidic PCR system. The CCHF virus and *Theileria s*pp. were specifically investigated in their respective potential vectors. DNA of eleven pathogens from six genera was identified in Corsican ticks and among them seven were reported for first time in Corsican ticks: *B. miyamotoi, B. afzelii, R. helvetica, A. phagocytophilum*, *Bar. henselae*, *Ba*. *bigemina* and *Ba. ovis*. These results also confirmed the presence of four other TBPs in Corsica: *R. aeschlimannii, R. slovaca*, *A. marginale*, and *T. equi*. Many of the pathogens found in this survey, as well as among those found by others, are mostly asymptomatic or benign for animals, highlighting domestic and wild Corsican animals are probably an important epidemiological reservoir increasing the human exposure to these zoonotic pathogens. These findings highlight the importance to look deeper into TBPs epidemiological situation in Corsica in the near future.

## ETHIC STATEMENT

The authors obtained the agreement of all the owners to collect ticks from their domestic animals. The cattle inspected were slaughtered for the human consumption and the wild boars collected were legally hunted during the hunting season. The collected deer, mouflons and birds were captured by PNRC and ONCF and they were all released. This study was approved by the veterinary Institutes (DDCSPP of Corse-du-Sud and Haute-Corse) and the French Ministry of Agriculture, General Directorate for Food (DGAl).

## ACKNOWLEDGEMENTS

The authors are grateful to the staff of the slaughterhouse of Ponte-Leccia, Cuttoli and Porto-Vecchio for their help in collecting ticks from cattle and to ONCFS, PNRC, hunters (especially Oscar Maestrini, INRA, Corte, France) and practicing veterinarians for collecting ticks respectively from mouflons, deer (and birds), wild boars and domestic carnivores.

